# Disruption of P2Y2 signaling promotes breast tumor cell dissemination by reducing ATP-dependent calcium elevation and actin localization to cell junctions

**DOI:** 10.1101/2023.03.31.533191

**Authors:** Makenzy L. Mull, Stephen J.P. Pratt, Keyata N. Thompson, David A. Annis, Abanoub A. Gad, Rachel M. Lee, Katarina T. Chang, Megan B. Stemberger, Julia A. Ju, Darin E. Gilchrist, Liron Boyman, Michele I. Vitolo, W. Jonathan Lederer, Stuart S. Martin

## Abstract

The tumor microenvironment and wound healing after injury both contain extremely high concentrations of the extracellular signaling molecule, adenosine triphosphate (ATP) compared to normal tissue. P2Y2 receptor, an ATP-activated purinergic receptor, is typically associated with pulmonary, endothelial, and neurological cell signaling. Here we report its role and importance in breast epithelial cell signaling and how it is altered in metastatic breast cancer. In response to ATP activation, P2Y2 receptor signaling causes an increase of intracellular Ca^2+^ in non-tumorigenic breast epithelial cells, while their tumorigenic and metastatic counterparts have significantly reduced Ca^2+^ responses. The non-tumorigenic cells respond to increased Ca^2+^ with actin polymerization and localization to cell edges, while the metastatic cells remained unaffected. The increase in intracellular Ca^2+^ after ATP stimulation was blunted using a P2Y2 antagonist, which also prevented actin mobilization and caused cell dissemination from spheroids in non-tumorigenic breast epithelial cells. Furthermore, the lack of Ca^2+^ concentration changes and actin mobilization in the metastatic breast cancer cells could be due to reduced P2Y2 expression, which correlates with poorer overall survival in breast cancer patients. This study elucidates rapid changes that occur after elevated intracellular Ca^2+^ in breast epithelial cells and how metastatic cancer cells have adapted to evade this cellular response.

## BACKGROUND

Wound healing and the tumor microenvironment (TME) have many parallel molecular markers, gene expression patterns, and phenotypes including, inflammation, proliferation, and migration.^1–3^ Moreover, genomic studies have shown that invasive tumors and wound repair mechanisms have similar gene expression patterns and display similarities between microenvironments.^3,4^ The TME consists of intra- and extracellular communication between non-malignant, resident and recruited cells, and the transformed tumor cells.^5,6^ One factor present at high, persistent levels within the TME is adenosine triphosphate (ATP), released by tumor and stromal cells in a regulated or non-regulated manner.^7–9^ Pellegatti *et al* found that ATP concentration ranged in the hundreds of micromolar in the tumor interstitium, while normal tissue has undetectable levels of extracellular ATP.^8^ This extracellular ATP, well-studied in neurotransmission, is an ideal messenger with high sensitivity due to low extracellular physiological concentrations, a rapid off-switch to avoid overstimulation or desensitization from a multitude of extracellular nucleotidases, and its ability to easily diffuse through aqueous tissues.^7,10^ These signaling and TME characteristics, make it critical to investigate the role of extracellular ATP in tumor signaling, immunosuppression, growth, migration, and invasion.

Activated by extracellular ATP, uridine triphosphate (UTP), adenosine diphosphate (ADP), and uridine diphosphate (UDP), the P2 receptors play a diverse role in neurotransmission, developmental biology, cardiac and pulmonary function, the apoptotic cascade, metastasis, as well as many other bodily functions.^11^ P2 receptors are divided into two families, P2X, a ligand-gated ionotropic channel, and P2Y, a metabotropic G-protein coupled receptor (GPCR), this paper will focus on the latter.^11^ ATP signaling via P2Y2 receptors has been shown in a few cancer studies, like prostate, liver, pancreatic, colorectal, and nasopharyngeal, but has remained more elusive in breast cancer research.^12–17^ A previous study showed altered purinergic receptor-Ca^2+^ signaling in endothelial growth factor (EGF)-induced epithelial mesenchymal transition (EMT) and later in hypoxia-induced EMT in breast cancer cells.^18,19^ While these previous studies have found correlation between purinergic signaling and cancer, most studies are using phenotypic assays based in cancer biology, which probe for slow cellular responses may take hours and days to respond. In contrast, we investigate rapid phenotypes occurring within seconds to minutes after purinergic receptor activation via ATP and how responses might vary in breast cancer compared to non-tumorigenic cells. Interestingly, purinergic signaling, specifically P2Y2 receptor, has been understudied in breast cancer and remains an interesting target to further investigate for therapy.^20–22^

Current breast cancer treatments include surgery, systemic therapy based on subtype, and chemotherapy, but metastatic disease remains nearly incurable.^23^ Approximately 1 in 8 women will be diagnosed with breast cancer in her lifetime, making female breast cancer the most diagnosed new case of cancer and the second leading cause of cancer death.^24^ Breast cancer can be organized into 3 subtypes based on gene amplification of epidermal growth factor 2 (ERBB2) and expression of estrogen or progesterone receptors.^23^ While available systemic therapies target *HER2* or hormone receptor positive subtypes, triple negative breast cancer (TNBC) lacks all three markers, and therefore, is more metastatic and most challenging to treat. 5-year survival rate can also vary based on the stage of breast cancer at diagnosis. There is an 86-96% 5-year survival rate for regional and localized breast cancers, while distant breast cancers have only a 29% 5-year survival rate.^24^ Another marker of poor survival in cancer patients is *KRAS*, a proto-oncogene and member of the RAS family of small GTPases. Only a single amino acid substitution is needed for an activating mutation at codon 12, 13, or 61.^25^ Activating *KRAS* mutations have been associated with various human cancers, including lung adenocarcinoma, colorectal, and pancreatic carcinomas.^26^ In breast cancer patients, *KRAS* is not the most common genetic mutation but it is present in around 30% of TNBCs.^27^ Previous work in our lab has led to the development of MCF10A cells with step-wise mutations of a *PTEN* double knockout and a *KRAS* activating mutation (10A-PTEN^-/-^KRas), that progressively push the cells towards tumorigenic transformation and metastasis.^28^ With all of the variations of breast cancer subtype and stage at the time of diagnosis, treatment plans and prognosis can range drastically for patients. Further research is needed to better target TNBCs and metastatic breast cancer.

Some research has linked P2Y2 receptor activation to increased migration via downstream modulators of the Ras pathway, like MEK-ERK in MCF7 breast cancer cells.^29^ Our lab previously showed that a scratch assay, similar to a wound edge, on breast epithelial cells transiently released ATP extracellularly, which activated P2Y2 receptor in neighboring cells and yielded a signaling cascade which increased intracellular calcium (Ca^2+^).^30^ Other research has linked increased intracellular Ca^2+^ and actin polymerization around the nucleus and endoplasmic reticulum in mammalian epithelial, mesenchymal, endothelial, and immune cells.^31^ Lasting only minutes before the actin cortex reformed, this rapid response was termed calcium-mediated actin reset (CaAR).^31^ Elevated intracellular Ca^2+^ concentrations also affect T lymphocyte activation, which is mediated by T cell receptor (TCR) clustering and dependent upon actin cortex interactions and movement to the cell surface.^32^ When there is an increase in intracellular Ca^2+^ leading to actin polymerization and increased interactions with TCR, reducing its mobility to the cell surface.^32^ Furthermore, Ca^2+^ fluxes and remodeling of the actin cytoskeleton have been evolutionarily conserved, and shown to play a role in *Drosophila* epithelial wound closure.^33^ Whether ATP-P2Y2-Ca^2+^ signaling in breast cancer has a similar molecular and functional mechanism regarding actin polymerization is still unknown.

It is well known that high extracellular concentrations of ATP are found in the TME, similar to a wound edge, and here we report that ATP alone is sufficient to stimulate an intracellular Ca^2+^ response in multiple breast epithelial and metastatic cancer cell lines. We demonstrate that ATP-activated P2Y2 receptors lead to a substantial increase in intracellular Ca^2+^ and a rapid polymerization of the actin network in non-tumorigenic breast epithelial cells. In contrast, metastatic breast cancer cells have an attenuated response to ATP with only a modest increase in intracellular Ca^2+^ and lack on actin re-localization. We also show that metastatic breast cancer cells have decreased expression of P2Y2 receptors, which could lead to their blunted Ca^2+^ response. Similarly, a chemical inhibitor of P2Y2 is able to recapitulate the insensitivity to ATP in the responsive non-malignant breast epithelial cells. Finally, we show that spheroids made of non-tumorigenic breast epithelial cells will disseminate in a 3-D collagen matrix when treated daily with P2Y2 inhibitor compared to control cells. We conclude that a non-malignant cell in the TME or at the wound edge would respond to ATP by increasing Ca^2+^ that would cause the actin cortex to stabilize and hold onto neighboring cells, while in metastatic and tumorigenic cells this ATP wound signal is muted, their actin cortex is unchanged, and migration can proceed unchecked.

## METHODS

### Reagents

P2Y2 receptor antibody was purchased from Abcam Inc (Cat#ab272891, Cambridge, UK). Phalloidin conjugated to Alexa Fluor Plus 647 (A30107) was purchased from Invitrogen-Thermo Fisher, Waltham, MA, USA. Wheat germ agglutinin (WGA)-Alexa 488 conjugate (Invitrogen, Cat#W11261). ATP (Sigma, Cat#A2383, Burlington, MA, USA) was reconstituted in double distilled H_2_O at pH 7.2 buffered with 20mM HEPES (Gibco-Thermo Fisher, Cat#15630-080, Waltham, MA, USA) and used at 10mM. Fluo-4 AM (Life Technologies-Thermo Fisher, Cat#F14201, Waltham, MA, USA) was made to working stock of 1mM according to manufacturer’s instructions and used at 4mM. Fluo-4 Direct Calcium Assay Kit, Starter Pack was used per manufacturer’s instructions (Invitrogen, Cat#F10471). AR-C 118925XX, referred to as P2Y2i, was reconstituted in DMSO according to manufacturer’s instructions (Biotechne-Tocris, Cat#4890, Minneapolis, MN, USA) and used at 10µM. Dimethyl sulfoxide (DMSO) was used as a control and dilutant for P2Y2i, purchased from Sigma-Aldrich (Cat#276855, Burlington, MA, USA).

### Cell Culture

Human MCF10A breast epithelial cells and human breast MDA-MB-231 cancer cells were obtained from the American Type Culture Collection (ATCC, Manassas, VA, USA) and were authenticated by short tandem repeat (STR) analysis. The creation of 10A-PTEN-/-KRas cells and their tumorigenicity have been previously described.^28^ MDA-MB-231 cells were also obtained from ATCC and authenticated by STR analysis, The MDA-MB-231 cells were maintained at 37°C, 5% CO_2_ and 95% humidity in Dulbecco’s Modified Eagle Medium (Corning, Cat#10-017-CV, Corning, NY, USA) supplemented with 10% FBS (Atlantic Biologicals, Cat#S11150H, Miami, FL, USA) and 1% penicillin-streptomycin (Gemini Bioproducts, Cat#400-109, West Sacramento, CA, USA). MCF10A and derivative cell lines were maintained in DMEM/F-12 Media (Invitrogen, Cat#10565-018) supplemented with 5% Horse Serum (Invitrogen, Cat#26050-088), 1% Pen/Strep, Recombinant Human EGF (Invitrogen, Cat#PHG0313, 100 µg/500 mL), Hydrocortisone (Sigma, Cat#H-0135, 0.5 µg/mL), Cholera Toxin (Sigma, Cat#C-8052, 100 ng/mL), and Insulin (Sigma, Cat#I-9278, 10 µg/mL). To maintain stocks, cells were passaged using a brief wash in PBS (Quality biological, Cat#114-058-101, Gaithersburg, MD, USA) followed by incubation with 0.25% Trypsin and 2.21mM EDTA (Corning, Cat#25-053-CI) at 37°C, 5% CO_2_ and 95% humidity.

### Live Imaging

Cells were plated to ∼80% confluency 48 hours before use in an IBIDI 8-well chamber slide with IBITreat (200 µL/well) (Cat#80826, Fitchburg, WI, USA). Cells were loaded with a cytosolic calcium sensitive dye, 4µM Fluo-4 AM in Hanks Balanced Salt Solution containing calcium (HBSS+Ca^2+^, Gibco, Cat#14025-092) for 30 minutes and then washed out and imaged with HBSS+Ca^2+^. For some experimental conditions cells were also treated with a selective and competitive P2Y2 antagonist, AR-C 118925XX or P2Y2i, which was reconstituted to 10mM stock in DMSO and used at a 10μM working dilution. After dye loading and washing, dishes were incubated in 10μM of P2Y2i or control DMSO (0.1%) for 10 minutes prior to imaging, without washout. Different DMSO concentrations and H_2_O controls did not significantly affect calcium signaling.

Epifluorescent, time-lapsed images were captured with a Nikon Ti2E microscope. All images were captured using a 20x phase-contrast objective (NA = 20x/0.45) with equal exposure time (100ms), interval time (5sec), and total time (10min). After a 30sec baseline read, 10mM ATP or control (H_2_O) was added during imaging. ATP (Sigma, Cat#A2383) was reconstituted in double distilled H_2_O at pH 7.2 buffered with 20mM HEPES (Gibco, Cat#15630-080). Nikon NIS-Elements was used to measure total field-of view relative fluorescent units (RFUs) over time before transfer of raw data to Excel and then analysis in GraphPad Prism 9.0.

### Quantitative Plate Reader

Cells were plated to ∼80% confluency, 48hrs before experiment in 96-well black, clear-bottom plates (200 µL/well). Cells were loaded with Fluo-4 made via the manufacturer’s directions from the Fluo-4 Direct Calcium Assay Kit, Starter Pack (Invitrogen, Cat#F10471) for 30 mins at 37°C, 5% CO_2_. Cells were then allowed 30mins to equilibrate to room temperature inside the FlexStation III plate reader (Molecular Devices, San Jose, CA, USA). An initial 30sec of baseline measurement at Excitation 488nm and Emission 515nm, before 10µM ATP or control ddH_2_O was added at 31 secs. Flow rate was set at the lowest setting (∼16 µL/sec) and 50 µL of ATP was added to 100 µL of cells/media at a depth of 80 µL. Change in fluorescence was recorded for 330 secs after ATP addition. SoftMax Pro files were converted to Excel and then analyzed in GraphPad Prism 9.0.

### RNA Seq

RNA samples were harvested from cells using the RNeasy kit Mini (Qiagen, Cat#217004, Germantown, MD, USA). The concentration was determined using the NanoDrop ND-1000 Spectrophotometer. Samples were frozen at -80^o^C until processed at the University of Maryland Baltimore’s Institute for Genome Sciences. Raw data from read counts of an n=3.

### Western Blotting

Cells were lysed using a chilled solution of 1× RIPA containing protease inhibitor cocktail, and phosphatase inhibitor cocktail II (Millipore-Sigma, Cat#20-188). Cells were kept chilled and scraped from the dish using a cell scraper. Lysates were collected in 1.5 mL microcentrifuge tubes and vortexed every 10min for the next 30min. The tubes were centrifuged at 15,000 rcf for 10min, before supernatants were collected in new microcentrifuge tubes. The resulting total protein was quantified using the Bio-Rad DC Protein Assay Kit according to the manufacturer’s recommended protocol (Bio-Rad, Cat#5000111, Hercules, CA, USA). Samples and standards were read on a BioTek Synergy HT plate reader after a 15 min incubation period. Samples were diluted to a final concentration of 1 µg/µL, boiled at 95 °C for 10 min, before gel loading. An amount of 20 µg of total protein was added to each lane of a 1.5 mm × 10 well NuPAGE 4–12% Bis-Tris Gel (Invitrogen, Cat#NP0335BPX). Loaded gels were run using 1× NuPAGE MES SDS Running Buffer at 90 volts for 30 min, followed by 120 volts for 90 min. Gels were transferred to a PVDF membrane using the eBlot™ L1 Fast Wet Transfer System (GenScript, Piscataway, NJ, USA). Membranes were blocked in 5% non-fat dry milk in 1× TBST rocking for 1hr at room temperature. Primary antibodies were used per manufacturer’s instructions and added to 5% non-fat dry milk in 1× TBST shaking overnight at 4 °C. Blots were washed 3 times with 1× TBST shaking for 10 min each. HRP-conjugated secondary antibody was diluted at 1:5000 in 5% non-fat dry milk in 1× TBST and incubated for 2hrs at room temperature with gentle rocking. Blots were washed 3 times with 1× TBST shaking for 10 min each. Electrochemiluminescence reagent (Amersham, Cat#RPN2232, Amersham, UK) was added to blots for 1min before capturing images on the iBright imager (Thermo Fisher). Densitometry was performed in ImageJ with Fiji (RRID:SCR_003070, RRID:SCR_002285).

### Immunofluorescence

Cells were plated to ∼80% confluency 48 hours before use in an IBIDI 8-well chamber slide with IBITreat (200 µL/well). HBSS+Ca^2^ was added to all cells for 1hr and used as the control. If treating with P2Y2i, drug or DMSO (0.1%) were added to cells for 10 mins at 10µM in the HBSS+Ca^2+^. At the various time points, 10µM ATP or control ddH_2_O was added to the HBSS+Ca^2+^. Cells were fixed using 3.7% formaldehyde diluted in 1x phosphate buffered saline (PBS) for 10 min and henceforth protected from light. The fixed cells were washed twice with PBS for 5mins each before WGA-Alexa 488 conjugate (Invitrogen, Cat#W11261) staining following manufacturer’s protocol for labeling pre-fixed cells. Next, the cells were permeabilized using 0.25% Triton-X100 (USB, Cat#9002-93-1, Cleveland, OH, USA) in PBS for 10 min followed by 2 more washes of PBS. Phalloidin (Invitrogen, Cat#A30107) was diluted in PBS at 1:1000, for nuclear co-staining a 1:1000 dilution of Hoechst (Millipore-Sigma, Cat#33258) was added to the solution. Cells were stained for 1hr at room temperature with gentle rocking before being washed with PBS 3 times at 10 mins each. A final wash with ultra-pure double distilled water was performed before drying, fixing with Fluoromount-G (Invitrogen, Cat# 00-4958-02), and storing at 4^o^C. Fixed slides were imaged on an Olympus FV-1000 confocal microscope at 60X magnification (NA = 60x/1.42 oil).

### 3-Dimensional Cell Dissemination Assay

#### Cell Aggregation and Spheroid Formation (Days 0-3)

confluent cells were washed with PBS and treated with 0.25% Trypsin to detach cells. Cells were then counted and diluted to 1x10^6^ cells/mL based on total volumed needed (100µL per well) before centrifugation at 1,000 RPM for 5mins. Spheroid formation media was made in a 3:1 ratio of cell culture media:MethoCult H4100 (StemCell Technologies, Cat#04100, Cambridge, MA, USA), and the total volume was calculated 100µL per well. Cell pellets were resuspended in spheroid formation media and plated 100 µL per well in a round bottom low-attach 96-well plate (Corning, Cat#7007). The whole 96-well plate was then centrifuged to allow cells to aggregate at 1,400 RPM for 7mins, then rotating the plate for a repeated centrifugation. Cell aggregates are allowed to grow undisturbed for 72hrs to form spheroids at 37°C, 5% CO_2_ and 95% humidity.

#### Spheroid Embedment (Day 3)

collagen I solution (3mL total) was prepared on ice following the protocol from Doyle, A. D. Generation of 3D collagen gels with controlled, diverse architectures. *Curr Protoc Cell Biol* **72**, 10.20.1-10.20.16 (2016).^34^ Collagen Type I was purchased from Corning (Cat#354236) and prepared at 2mg/mL. Reconstitution buffer was prepared and NaOH was added to balance the pH, both following the protocol. Chilled cell culture media was used to make up the volume to 3mL. Each spheroid was gently transferred from the round bottom 96-well plate in a 1mL pipette tip to a petri dish for isolation from MethoCult H4100 mixture. 100µL of cold collagen I solution was added to a flat bottom low-attach 96-well plate, pre-warmed to 37°C on a heat block within the cell culture hood. The collagen solution was allowed to partially polymerize for 2mins, before collecting the spheroid in collagen solution and adding to the middle of the well. Spheroids were allowed to embed in collagen for at least 2mins at 37°C. These steps were repeated for each spheroid, then the whole plate was incubated for 30mins at 37°C, 5% CO_2_ and 95% humidity, to allow the collagen to fully polymerize. An additional 100µL of warm cell culture media was added on top of the gels. Spheroids and 3-D cell dissemination were imaged from Day 0 through Day 7 at 10X phase contrast (NA=10x/0.30), and image tiling (2x2) was implemented when needed on a Nikon Ti2E inverted microscope. When indicated, some experimental conditions were treated with P2Y2i at 10µM or control (DMSO 0.1%) in media and replenished every 24hrs. Images were analyzed with NIS-Elements AR (Nikon) software. Spheroid perimeter is traced, and area is measured in µm^2^, then change in area normalized to Day 0 is calculated in GraphPad Prism 9.0.

## Bioinformatics

The online cBioPortal (RRID:SCR_014555) database (https://www.cbioportal.org/) was used to test whether P2Y2 mRNA expression had an effect on disease free survival in two different breast cancer datasets. The online Kaplan–Meier Plotter (RRID:SCR_018753) database (https://kmplot.com/analysis/) was used to test whether P2Y2 mRNA expression had effects on breast cancer patient clinical outcome. The Gene symbol used was P2RY2.

### Statistics

Statistical analyses were conducted using two-tailed t-test, one-way ANOVA with Dunnett’s or Holm-Sidak multiple comparison post-tests, or two-way ANOVA with Geisser-Greenhouse correction and Tukey’s multiple comparisons test in GraphPad Prism 9.0 software; P < 0.05 was considered significant.

## RESULTS

### Extracellular ATP stimulates calcium signaling in normal and tumorigenic breast epithelial cells

Following up on previous work by the lab, we aimed to test adenosine triphosphate (ATP) as a signaling molecule for intracellular calcium across multiple breast epithelial cell lines.^30^ Since there are few studies that have focused on rapid calcium signaling in breast cancer, we wanted to measure signaling in tumorigenic cell lines as well. Time-lapsed images were captured with epifluorescence on a Nikon Ti2E microscope, and relative fluorescent units (RFUs) were measured in Nikon NIS-Elements software (Sup. Vid. 1-3). Representative images from those compressed videos were selected at specific time points: before and after ATP addition, 1 min, and 10 min (Fig. 1A). We observed that non-tumorigenic MCF10A cells have a rapid and significant intracellular Ca^2+^ response after 10mM ATP addition, indicated in green (Fig.1A). 10A-PTEN-/-KRas and MDA-MB-231 cells both responded with intracellular calcium increases after ATP stimulation, but both failed to reach the increase in RFUs compared to non-tumorigenic MCF10A (Fig. 1A). Whole-field change in fluorescence compared to baseline fluorescence (△F/F_0_) as well as peak RFU were both greater in MCF10A cells after ATP treatment, while the two metastatic breast tumor cell lines showed a comparatively lower induction of cytoplasmic calcium after ATP addition (Fig.1B-D). Only MCF10A cells had a significant difference between ATP-treatment and control peak RFU, as well as MCF10A + ATP compared to MDA-MB-231 + ATP (Fig. 1E). When comparing all conditions and cell lines to one another, the MCF10A cells had a significantly greater intracellular Ca^2+^ response than the two tumorigenic cell lines (Fig. 1E). In order to confirm our results, we used real-time quantitative plate reader measurements as an orthogonal approach. The increase in intracellular Ca^2+^ response was also seen in the MCF10A cells treated with ATP in the FlexStation III (Sup. Fig. 1A). The 10A-PTEN-/-KRas cells and MDA-MB-231 cells showed a much smaller intracellular Ca^2+^ response (Sup. Fig. 1B-C). Area under the curve (AUC) measurements were performed and a one-way ANOVA with multiple comparisons was used to calculate significance (Sup. Fig. 1D). Comparing all three cell types, we again saw a significant difference in Ca^2+^ response from the MCF10A cells, while there were no significant changes in the metastatic cell lines (Sup. Fig. 1D). Overall, these results illustrate that stimulating breast epithelial cells with ATP leads to a rapid Ca^2+^ response that is blunted in aggressive tumor cells.

**Figure 1.**
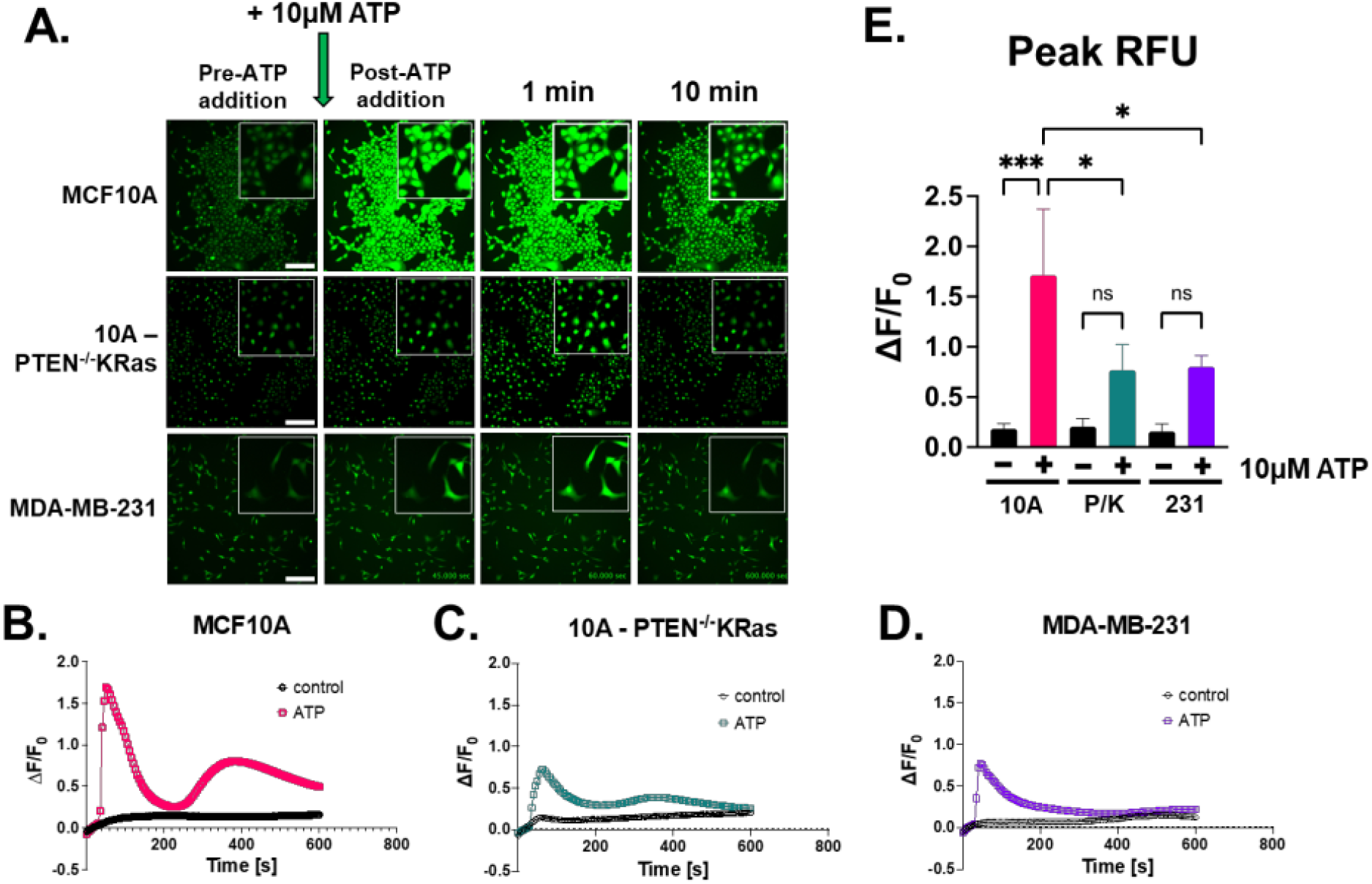
ATP as a pseudo-mechanical stimulus causes intracellular calcium to increase in breast epithelial cells. **A)** 20x epi-fluorescent time-lapsed images of MCF10A, 10A-PTEN^-/-^ KRas, and MDA-MB-231 cells loaded with Fluo-4 AM to label calcium. 10µM ATP was added after a 30 sec baseline read. Representative images of pre-ATP addition, post-addition, 1 min, and 10 min were taken (Scale bar = 200 µm) and insets at 4x zoom. **B)** Graph of △F/F_0_ from whole field of view showed a rapid Ca^2+^ response in MCF10A cells treated with 10µM ATP. **C)** 10A-PTEN-/-KRas (P/K) cells saw a suppressed intracellular Ca^2+^ response. **D)** Similarly, MDA-MB-231 cells also had an inhibited Ca^2+^ response after ATP stimulation. **E)** Peak RFU with one-way ANOVA and Sidak’s multiple comparisons test were performed on graphs from each cell line with SD (***, P<0.001, *, P<0.05) (all graphs n=3).

### P2Y2 receptor expression is lower at the mRNA and protein levels in tumorigenic cells compared to MCF10As

After showing by multiple measurements that the non-tumorigenic MCF10A cells increased intracellular Ca^2+^ compared to metastatic cell lines, we wanted to investigate a potential mechanism responsible for this difference in our breast epithelial cell lines. We focused on our previous findings that extracellular ATP signals are conveyed through the purinergic receptor, P2Y2, in breast epithelial tissue.^30^ RNA sequencing data showed that P2Y2 expression is reduced nearly 8-fold in the mutant 10A-PTEN-/-KRas cells as well as the 10A-KRas cell line (Fig. 2A). The metastatic MDA-MB-231 and MDA-MB-436 cells from ATCC also showed a large fold-change in decreased P2Y2 mRNA expression compared to MCF10A cells (Fig. 2A). After discovering the large decrease in P2Y2 mRNA, we examined protein expression in seven different tumorigenic and non-tumorigenic breast epithelial cell lines. P2Y2 protein expression appeared to be down in 10A-PTEN-/-KRas and MDA-MB-436 cells but was significantly decreased in MDA-MB-231 cells compared to MCF10A cells (Fig. 2B and Sup. Fig. 2). Densitometry shows that there is a significant reduction in P2Y2 protein expression in MDA-MB-231 cells. While there was also a decrease in protein expression in the 10A-PTEN-/-KRas and MDA-MB-436 cells it did not reach statistical significance (Fig. 2C and Sup. Fig. 2). These data show that there is a significant decrease in P2Y2 expression at the mRNA and protein level in metastatic MDA-MB-231 cells.

**Figure 2.**
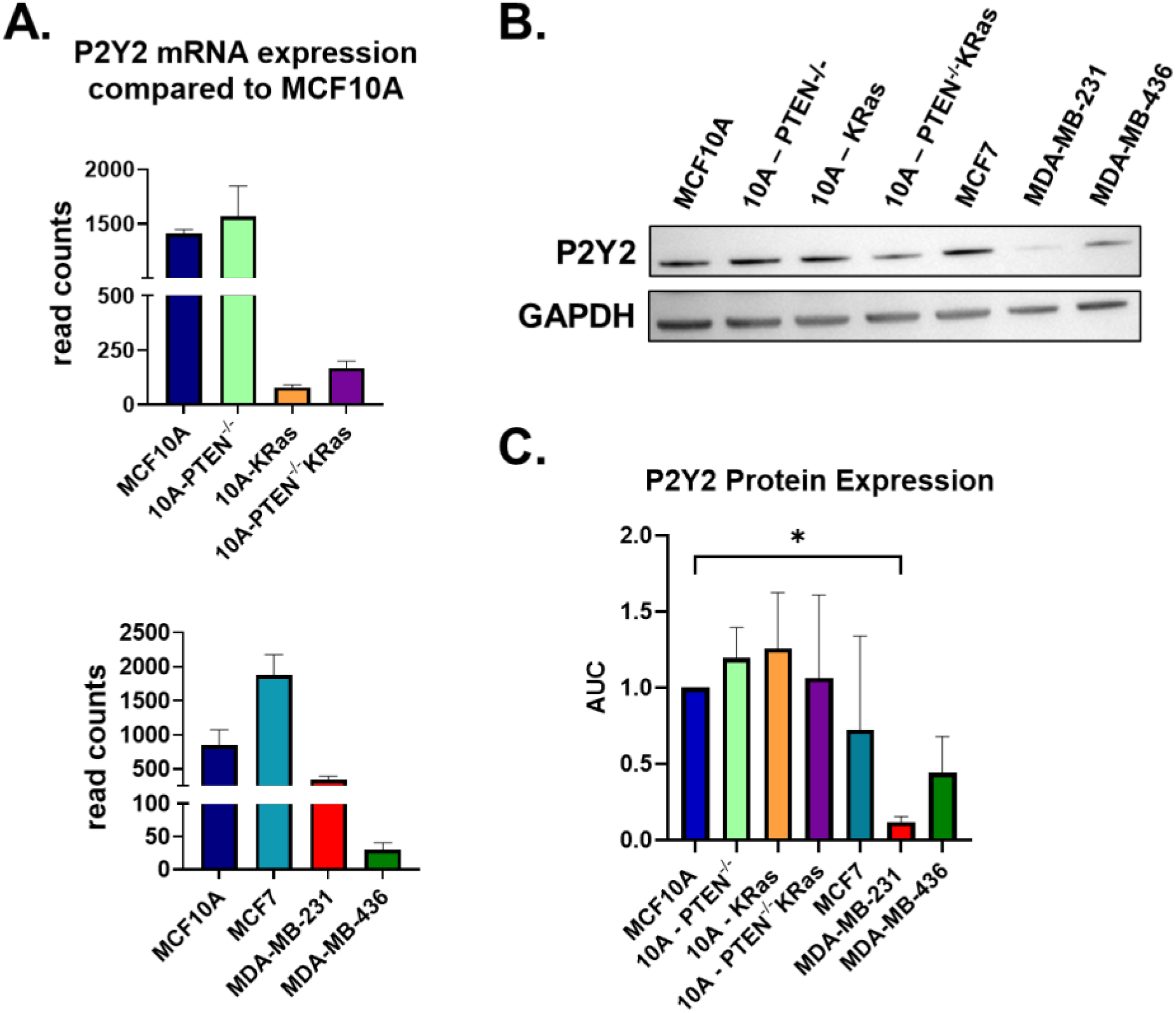
P2Y2 Expression is lower at the mRNA and protein level in mutant 10A-PTEN-/-KRas and TNBC cell lines compared to MCF10A. **A)** mRNA sequencing results show lower expression of P2Y2 receptor in 10A-KRas, 10A-PTEN-/-KRas, MDA-MB-231, and MDA-MB-436 cells in comparison to MCF10A control. **B)** Representative western blot for P2Y2 (42kDa) receptor across seven different cell lines show a decrease in P2Y2 expression in the metastatic cancer cell lines with GAPDH (37kDa) loading control. **C)** There were decreases in protein expression for the metastatic breast cancer cell lines, although the only significant difference was in MDA-MB-231 cells. Quantification of western blots with P2Y2 expression normalized to GAPDH then compared to MCF10A with SD (*, P<0.05) (n=3) using one-way ANOVA with Dunnett’s multiple comparisons test.

### ATP-induced intracellular Ca^2+^ response is inhibited by P2Y2 antagonist in breast epithelial cells

Previous work in our lab showed that AR-C118925XX, a P2Y2 antagonist that will be referred to as P2Y2i, was able to block a mechanically-induced calcium response in MCF10A and MDA-MB-231 cells after scratch assays.^30^ To test if P2Y2 receptors are involved in the differential response to ATP in mammary epithelial and breast tumor cells, we examined the effect of P2Y2i on ATP-stimulated Ca^2+^ elevation. Cells were pre-treated with 10µM P2Y2i for 10 mins after loading with Fluo-4 AM and time-lapsed imaging was performed similar to Figure 1 for all three cell lines. Figure 3A shows representative images from all three cells lines pre- and post-ATP addition, 1 min, and 10 min post-stimulation. Total field-of-view RFUs were analyzed in Nikon NIS-Elements, and graphed as △F/F_0_ control, ATP, or P2Y2i + ATP (Fig. 3B-D). In Figure 3B, P2Y2i significantly suppresses the ATP response in MCF10A cells, while breast tumor cells (Fig. 3C, D) are inhibited almost completely, similar to the control. Measuring peak RFU with one-way ANOVA statistical test, we found no significant differences in intracellular Ca^2+^ changes over time among the three cell lines treated with P2Y2i compared to control (Fig. 3E), indicating a complete suppression of the ATP-induced intracellular Ca^2+^ with P2Y2i. These results demonstrate that MCF10A cells, when treated with an antagonist for P2Y2 receptor and then stimulated with ATP, recapitulate a similar blunted Ca^2+^ response to the metastatic breast cancer cells.

**Figure 3.**
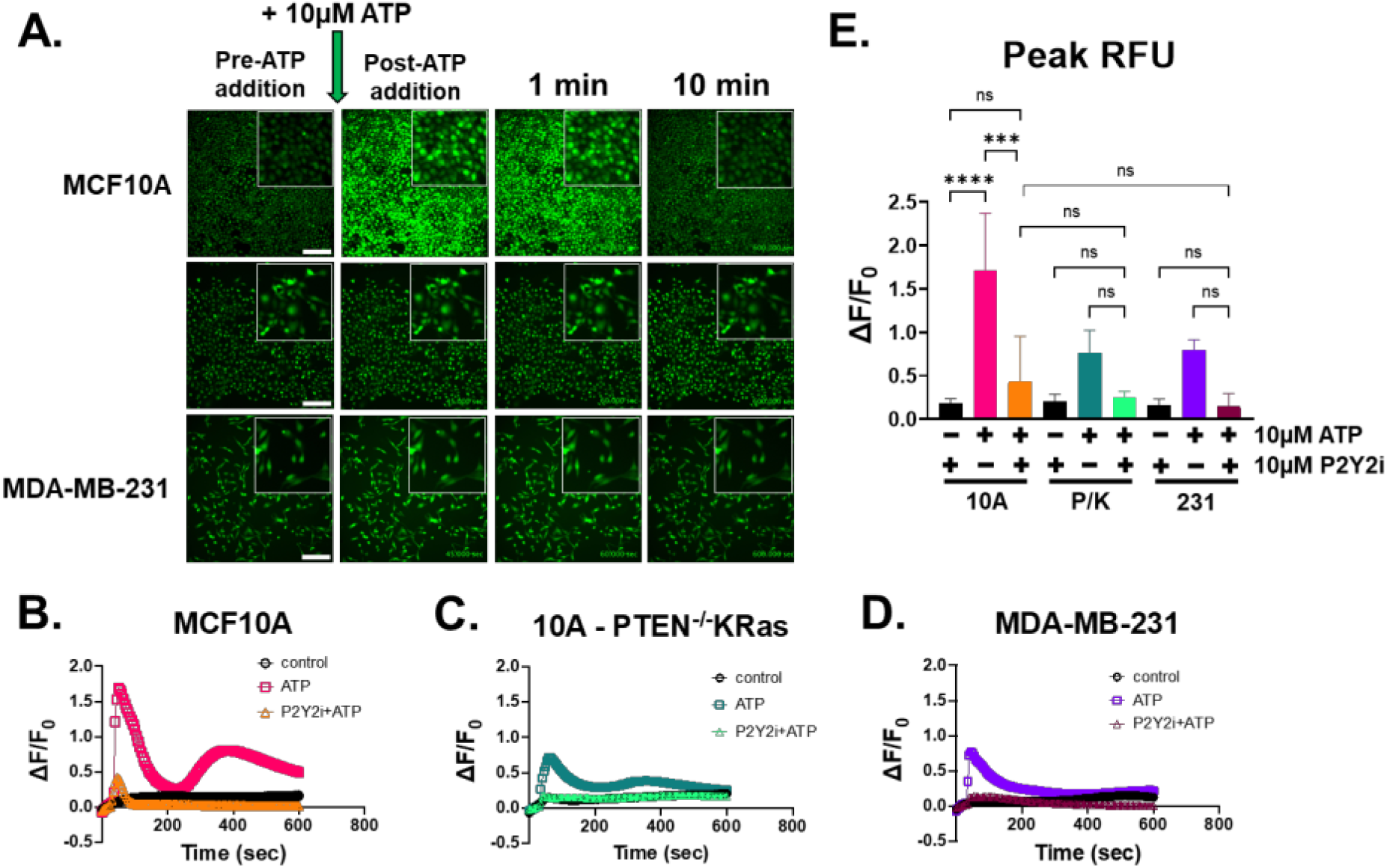
P2Y2 antagonist, AR-C118925XX, interrupts the calcium signal stimulated by ATP. **A)** Epi-fluorescent images of MCF10A, 10A-PTEN-/-KRas, and MDA-MB-231 cells loaded with Fluo-4 and treated with 10µM P2Y2i for 10 mins before imaging. Representative frames of pre-ATP addition, post-addition, 1 min, 5 min, and 10 min were taken from time-lapsed images (scale bar = 200 µm) and insets at 4x zoom. **B)** Graph of △F/F_0_ from whole field of view showed a suppressed Ca^2+^ response in MCF10A cells treated with 10µM P2Y2i before ATP stimulation compared to ATP with no antagonist. **C)** 10A-PTEN-/-KRas cells saw no significant change from control with P2Y2i treatment. **D)** MDA-MB-231 cells also had no significant Ca^2+^ changes with P2Y2i treatment. **E)** Peak RFU with one-way ANOVA and Sidak’s multiple comparisons were performed on graphs from each cell line with SD (****, P<0.0001) (n=3).

### Rapid changes in the actin cortex occur after ATP-induced calcium increases in breast epithelial cells

We have now shown that ATP via P2Y2 receptor can induce an intracellular Ca^2+^ response in breast epithelial cells, while tumorigenic cells have a significantly lower response compared to MCF10A cells. What remained unknown was the effect of that Ca^2+^ signal on cell behavior that could impact responses to stress, wound healing, or elevated ATP levels in the tumor microenvironment. It is well known that actin and Ca^2+^ work closely together, which led us to explore changes in the actin network after ATP treatment.^31^ We found that after treatment with 10µM ATP, MCF10A cells demonstrated a rapid actin reorganization that localizes at cell edges and junctions within 10 mins (Fig. 4A, top panel). The same actin polymerization and localization was not seen in the 10A-PTEN-/-KRas or MDA-MB-231 cells, supporting the idea that tumorigenic cells might not be responsive to normal wounding signals (Fig. 4B, C, top panels). Similar to our results with P2Y2i in Figure 3, we also show that P2Y2i prevents actin re-localization and polymerization after ATP stimulation in non-tumorigenic breast epithelial cells (Fig. 4A, lower panel). The metastatic breast cancer cells show no major changes in actin after P2Y2i treatment and ATP stimulation (Fig. 4B, C, lower panels). WGA (wheat germ agglutinin) was used to label the whole cell membrane and Hoechst to stain the nucleus (Sup. Fig. 3-4). These data show that breast epithelial cells have changes in the actin cortex in response to rapid Ca^2+^ signaling after ATP stimulation, which can be inhibited with P2Y2i, while the metastatic breast cancer cells lack a response.

**Figure 4.**
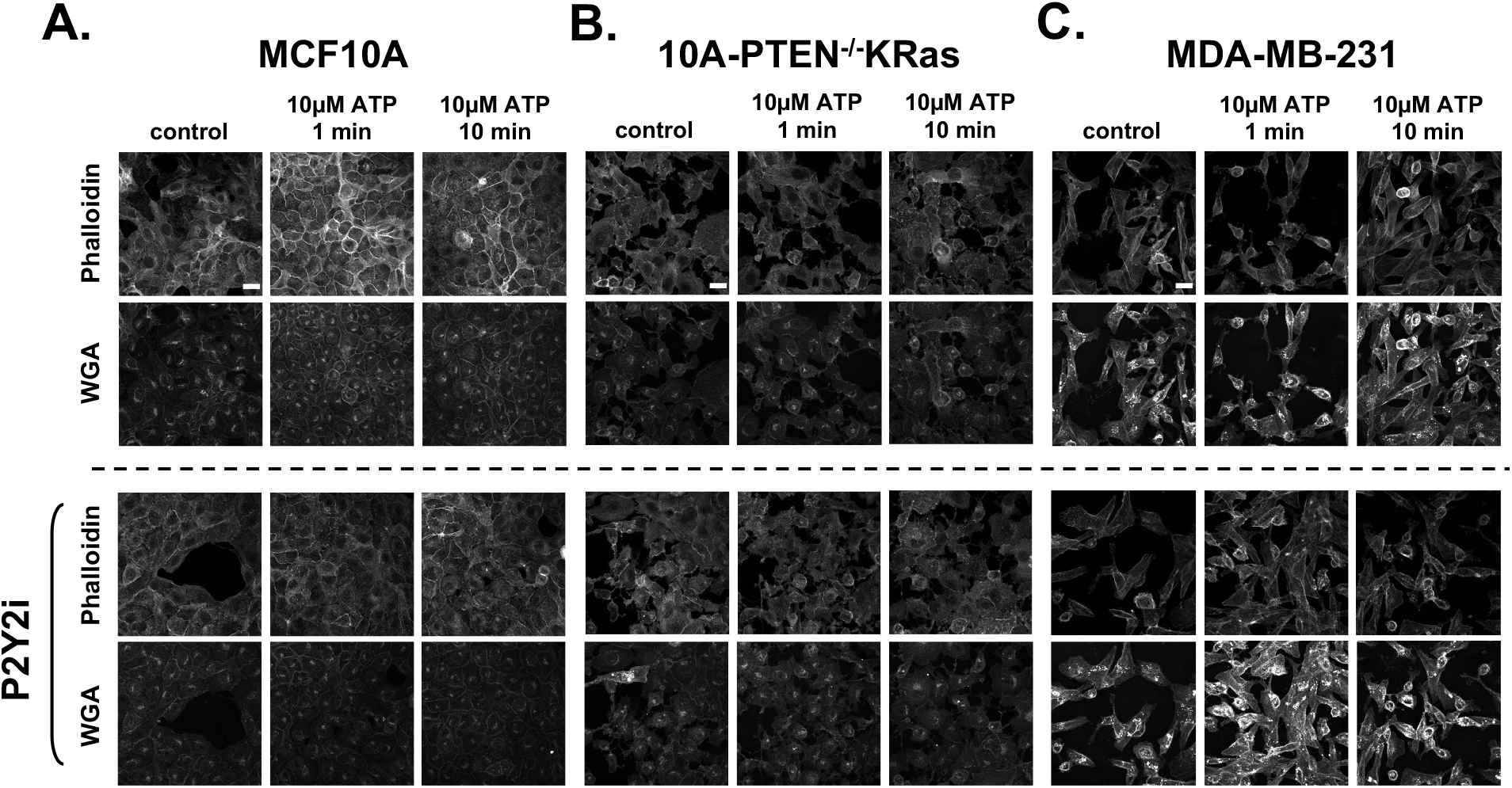
ATP-activated increase in calcium leads to rapid actin localizing around cellular junctions in non-tumorigenic breast epithelial cells. All cells were treated with control or 10µM ATP for 1 min, or 10 mins. **A)** Confocal images of MCF10A cells stained with phalloidin show changes in actin polymerization and localization to cellular edges and junctions after ATP stimulation. Although when using the P2Y2i, MCF10A cells show no changes in actin after ATP stimulation. **B)** Confocal images of our 10A-PTEN-/-KRas cells show no major changes in phalloidin staining after ATP addition with or without P2Y2i. **C)** Confocal images of MDA-MB-231 cells also show no changes in actin with control or P2Y2i and ATP stimulation. Confocal NA: 60x/1.42 oil. All images stained with Alexa conjugated Phalloidin (nm=647) and WGA-Alexa (nm=488) to stain cell membranes. Representative images were taken from a n=3. Scale bar=25µm.

### P2Y2i treatment increases cell dissemination and spheroid size in non-tumorigenic breast epithelial cells

Given the ATP-dependent actin reorganization at cell junctions and disruption of this response by P2Y2i (Fig. 4), we examined long-term impacts on epithelial cell dissemination from spheroids. Using an expedited method, we formed spheroids from homotypic cell aggregates in low-attach plates before transferring the spheroids into a 3-D collagen matrix and monitored growth and dissemination. We found that the control treated MCF10A cells had minimal invasion into the collagen matrix over several days (Fig. 5A top panel and Sup. Fig. 5). Oppositely, the MCF10A cells treated with P2Y2i were able to invade into the matrix and disseminate away from the spheroid, increasing the overall area (Fig. 5A bottom panel and Sup. Fig. 5). Images taken over the course of 7 days were quantified using whole spheroid area and the change in area from Day 0 (Fig. 4B). We show significant increases in whole area and change in area when cells are treated with P2Y2i compared to control (Fig. B). This could be due to the fact that pharmacological inhibition of P2Y2 also blunts intracellular Ca^2+^ increases and prevents rapid actin response, allowing cells to ignore these mechanical signals, similar to metastatic cells with lower expression of P2Y2 receptor. Therefore, the normally non-tumorigenic cells are insensitive to mechanical signals when treated with P2Y2i and will continue to migrate outward. We were also able to recapitulate this phenotype in MCF7 cells, which are tumorigenic but not normally metastatic *in vivo* (Sup. Fig. 6). Overall, these data show that P2Y2 inhibition is sufficient to increase 3-D cell dissemination in non-tumorigenic breast epithelial cells, while further establishing the role of P2Y2 receptor in breast epithelial cell signaling and sensing the local environment.

**Figure 5.**
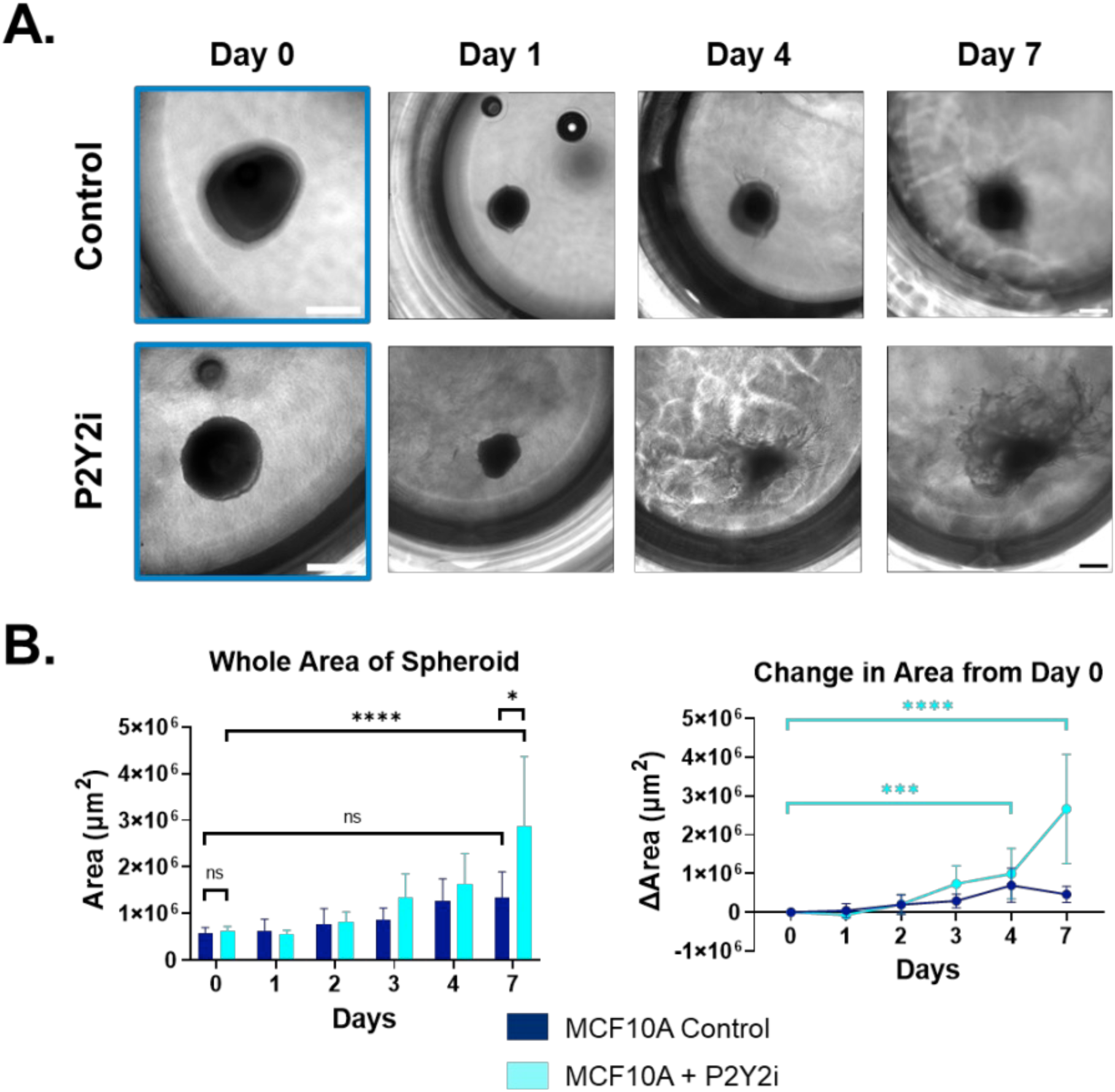
3-D Cell dissemination of non-tumorigenic MCF10A cells treated with P2Y2i. Spheroids formed for 72hrs were embedded into collagen solution and imaged every day for 7 days. **A)** Representative phase contrast images taken at 10x on Day 0 (blue outline) or 10x stitched (2x2) of spheroids treated with control or P2Y2i on Days 1, 4, and 7. Blue outlines represent 10x images unstitched (white or black scale bar=500µm). **B)** Graphs showing whole area of spheroid (left) and change in area (Δµm^2^) from day 0 through day 7 (right) with SD, and two-way ANOVA with Tukey’s multiple comparisons post-test performed to calculate significance (****, P<0.0001, *, P<0.05) (representative images from n=3).

**Figure 6.**
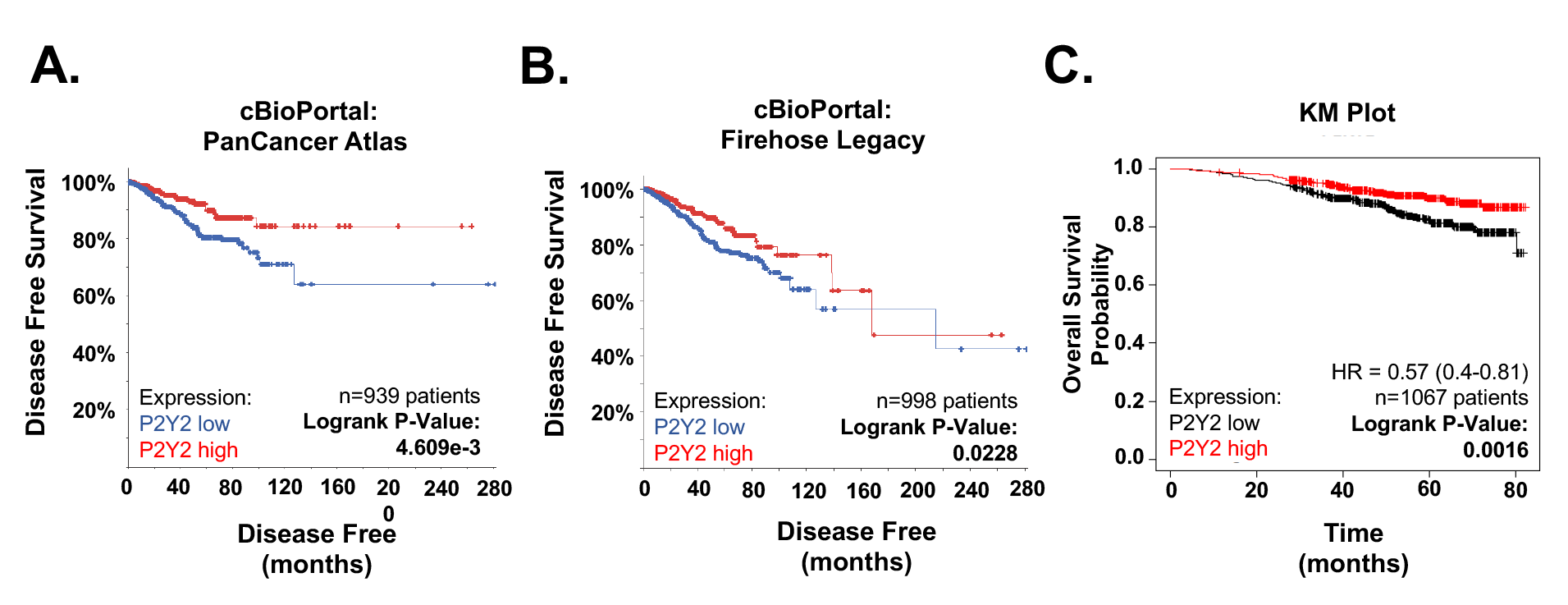
Three different survival curves for breast cancer patients based on expression of P2Y2 receptor. **A)** Data from cBioPortal: PanCancer Atlas patients with breast cancer show patient tumors with lower P2Y2 mRNA expression have poorer disease-free survival (P=4.609e-3). **B)** Data from cBioPortal: Firehose Legacy patients with breast cancer and low P2Y2 mRNA expression also have a smaller percentage of disease-free survival (P=0.0228). **C)** Kaplan-Meier Plot from KM Plot shows that low expression of P2Y2 receptor is correlated with poorer probability of overall survival (P=0.0016).

### Low P2Y2 mRNA expression is correlated with decreased survival and disease-free progression in breast cancer patients

To further support our findings, we used bioinformatic tools to explore mRNA expression versus patient outcomes. Gene expression analysis showed a strong correlation with poor survival and low disease-free outcomes in patients with reduced P2Y2 mRNA expressing tumors. Two cBioportal breast cancer datasets, the Invasive Breast Carcinoma (TCGA, PanCancer Atlas) and Firehose Legacy, which also includes invasive breast carcinoma samples, both showed significant correlation with low P2Y2 expression and smaller percent chance of disease-free survival (Fig. 6A,B).^35,36^ The independent KM Plot database also demonstrated that breast cancer patients of all subtypes, had reduced overall survival probability with tumors that had reduced P2Y2 receptor expression (Fig. 6C).^37^ Additionally, this clinical data help support our findings that metastatic breast cancer cells with less P2Y2 receptor expression have a limited Ca^2+^ response after ATP-activation.

## DISCUSSION

The tumor microenvironment (TME) is composed of many different cell types, and the role of non-malignant cells play a large factor in promoting tumors.^6^ ATP is released into the TME from multiple cell types and has been found in extremely high concentrations in the extracellular tumor space.^8,38^ ATP concentrations in the tumor interstitium have been found in the hundreds of micromolar, compared to normally undetectable levels in a non-malignant tissue.^8^ Since it is well known that ATP is an activator of purinergic receptor signaling, multiple studies have investigated the role of purinergic receptors, such as the P2Y2 receptor, in wound healing. Boucher *et al*. found that P2Y2 receptors play a key role in the response to corneal epithelial cell injury and repairs.^21^ As our lab previously observed in breast epithelial cells, mechanical damage to the corneal epithelial cells caused a release of ATP and a stimulation of purinergic receptors (P2Y2), which subsequently lead to a Ca^2+^ response.^21,30^ We also saw that the Ca^2+^ signal could be blocked if the cells were pretreated with apyrase, while Boucher *et al*. found that the purinergic signaling also lead to phosphorylation of ERK.^21,30^ Another group found that pharmacological inhibition of P2Y2 receptor, phospholipase C, and inositol 1,4,5-triphosphate receptor decreased wound closure in human keratinocytes.^39^ A previously unknown mechanism was shown by a group in 2013, indicating endothelial P2Y2 receptor involvement in cancer cell extravasation via activated platelet release of ATP.^40^ Since wound healing has many similarities to the TME, we pursued the high levels of ATP and have shown sufficiency of ATP alone to stimulate intracellular Ca^2+^ in breast epithelial cells (Fig. 1).

Purinergic receptor signaling in breast cancer is a unique niche, where a few groups have found involvement of ATP-activated P2Y2 receptor and relatively long-term cancer phenotypes that occur over hours to days, like epithelial-to-mesenchymal transition (EMT), migration, and invasion.^18,20,29,41,42^ Davis *et al*. found that initiation of EMT using endothelial growth factor in MDA-MB-436 breast cancer cells caused an increase in vimentin at 24 hours.^18^ Using RT-PCR, they found that P2X5 mRNA expression was increased in their cells and also found changes in Ca^2+^ signaling via ATP stimulation with increased vimentin.^18^ Parallel to that, here, we have identified differences in intracellular Ca^2+^ response after ATP-activated P2Y2 receptor signaling in metastatic breast cancer cells with oncogenic KRas mutations compared to non-malignant breast epithelial cells (Fig. 1). The main purpose of our study was to investigate the rapid phenotypes that occur within seconds to minutes and could lead to later changes in migration and invasion. Previous work led us to P2Y2 receptors, and while there is some variation in P2Y2 protein expression levels across breast epithelial cells and breast cancer lines, we saw a decrease in P2Y2 mRNA and protein in MDA-MB-231 cells compared to our MCF10A cell line (Fig. 2).^41,43^ The metastatic breast cancer cells might not respond to the external ATP signal due to their significantly decreased mRNA and protein expression of P2Y2 receptor, thereby suppressing Ca^2+^ flux within the cytoplasm.

Calcium concentration in the cytoplasm is normally around 100nM, while extracellular Ca^2+^ is much higher in the mM range.^44^ The field of calcium signaling in cancer is mainly focused on aberrant gene expression due to changes in Ca^2+^ concentrations or protein expression of receptors causing differences in Ca^2+^ concentrations.^44–46^ We, and others, have now shown differences in P2Y2 receptor expression, we have focused on the rapid Ca^2+^ changes and the phenotypes that occur afterwards. As previously studied, the role of Ca^2+^ and the actin cortex has been well established in muscle, bone, and cardiac research.^47^ The rapid polymerization and localization of actin in response to ATP-stimulated Ca^2+^ increase is a normal cell response to wounding. In parallel, we show metastatic tumorigenic cells with lower P2Y2 expression do not respond significantly to ATP, and therefore yield minimal flux of Ca^2+^ with no actin polymerization and reorganization at cell edges (Figs. 4 and 5). It would be advantageous for breast cancer cells to not respond or become desensitized to the high ATP concentrations in the TME and continue to migrate out of the primary tumor.

Since the P2Y2 receptor is part of the large family of purinergic receptors, there are a few drugs developed to target it or its family members. AR-C118925 is a potent and specific antagonist of the P2Y2 receptor.^48^ The drug was developed using UTP agonist as the base and is relatively new.^48^ Thus far, only a few P2Y2 drugs are being used clinically. Diquafosol is a P2Y2 receptor agonist that has been used as treatment for dry eye disease, since P2Y2 is expressed in the epithelium of the eye.^49^ Another antagonist for P2Y and P2X receptors is the non-selective drug Suramin.^50^ It has most commonly and consistently been used as a treatment for human sleeping sickness, but also has applications for viral disease like HIV and hepatitis, as well as studies looking at autism, snakebites, and cancer.^50^ Unfortunately, it is extremely indiscriminate and has a multitude of targets from DNA and RNA synthesis, sirtuins, protein kinases, tyrosine phosphatases, ATPases, GABA receptors, etc.^50^ There was even one clinical trial for Suramin and Paclitaxel in women with stage IIB-IV breast cancer from 2003 to 2015. The goal was to give the drugs in combination and to assess the best dose of Suramin that might increase the effectiveness of Paclitaxel by sensitizing the tumor cells.^51^ They found that Suramin at non-toxic doses in combination with Paclitaxel given weekly, was tolerated well by metastatic breast cancer patients, but the efficacy did not meet the criteria essential to rationalize further investigation.^51^ Purinergic receptors are such a large family that drugs must be specific to avoid as many off target effects as possible. A P2Y2 antagonist might not be the best choice in metastatic breast cancer cells with low expression of P2Y2 receptor compared to non-tumorigenic breast epithelial. Even in tumor cells with normal or elevated P2Y2 receptor expression, our results indicate treatment with a P2Y2i would be predicted to reduce actin-dependent reinforcement of cell junctions in the presence of ATP, which could inadvertently induce tumor cell dissemination. We show that non-tumorigenic MCF10A cells in control conditions do not significantly disseminate or increase spheroid size over 7 days, while MCF10A cells treated with P2Y2i significantly disseminate outward in a 3-D matrix (Fig. 5). There may be potential use for a P2Y2 agonist if that could play a role in sensitizing the cells to ATP signaling and increasing intracellular Ca^2+^ response.

## CONCLUSIONS

In summary, our results demonstrate that metastatic breast cancer cells have a suppressed intracellular Ca^2+^ response to ATP-stimulated P2Y2 receptor signaling compared to non-malignant breast epithelial cells. This insensitivity could be due to low expression of the receptor in metastatic breast cancer cells, supported by patient survival data, in which survival decreases with low P2Y2 expression. Further research into the rapid phenotypes occurring after P2Y2 activation could help elucidate better mechanisms for drug therapy and a deeper understanding of how cancerous cells evade typical mechanical signals.

## Supporting information

Combined Supplemental Figures and Legends

## LIST OF ABBREVIATIONS

ATP: Adenosine Triphosphate
ADP: Adenosine Diphosphate
UTP: Uridine Triphosphate
P2Y2: Purinergic Receptor P2Y2
TME: Tumor microenvironment
EGF: Epidermal growth factor
EMT: Epithelial to mesenchymal transition
TNBC: Triple negative breast cancer
Ca^2+^: Calcium
CaAR: Calcium-mediated actin reset
HBSS: Hanks’ Balanced Salt Solution
DMSO: Dimethyl sulfoxide
ddH_2_O: Double-distilled water
P2Y2i: Antagonist for P2Y2, AR-C 118925XX
WGA: Wheat germ agglutinin
RFU: Relative fluorescence units
ΔF/F_0_: Change in fluorescence over baseline fluorescence
AUC: Area under curve

## DECLARATIONS

### Ethics approval and consent for publication

Not applicabl

### Data Availability Statement

The datasets generated during and/or analyzed during the current study are available from the corresponding author on reasonable request. The datasets generated during and/or analyzed during the current study are available in the cBioPortal (RRID:SCR_014555) or KM Plot (RRID:SCR_018753) repository (see methods for further information). [https://www.cbioportal.org/] [https://kmplot.com/analysis/]

### Competing Interests

SSM is an employee of the VA Maryland Health Care System. The views reported in this paper do not reflect the views of the Department of Veterans Affairs or the United States Government. The PTEN^−/−^ cells are licensed by Horizon Discovery Ltd. (Cambridge, UK), MIV receives compensation from the sale of these cells. At the time of this publication, AAG is an employee of AbbVie inc.

### Funding

MLM was supported by 5T32GM008181-33 & 34 from the National Institute of General Medical Sciences. This work was supported in part by the METAvivor Foundation and grants to SSM from the US Department of Veterans Affairs (I01-BX002746) and the National Institutes of Health (R01-CA154624, R01-CA124704). MIV was supported by RSG-18-028-01. The Greenebaum Comprehensive Cancer Center is supported by P30-CA134274 and the Maryland Cigarette Restitution Fund.

### Authors’ Contributions

Conceptualization, MLM, SJPP, WJL, and SSM; methodology, MLM, SJPP, AAG, MBS, JAJ, LB, and SSM; validation, MLM; formal analysis, MLM, SJPP, AAG, JAJ, LB, and RML; investigation, MLM and SSM; resources, MLM, SJPP, KNT, DAA, JAJ, RML, AAG, MIV, LB, SSM ; data curation, MLM; data interpretation, MLM, SJPP, KNT, DAA, AAG, RML, JAJ, MBS, KTC, DEG, LB, MIV, and SSM; writing—original draft preparation, MLM and SSM; writing—review and editing, MLM, SJPP, KNT, DAA, AAG, JAJ, MBS, RML, KTC, DEG, MIV, LB, WJL, and SSM; supervision, KNT, MIV, LB, WJL, and SSM; project administration, MLM; funding acquisition, MLM and SSM. All authors have read and agreed to the final version of the manuscript.

## Notes

### Summary of Updates

New data added to figures and supplementary data. A new Figure 5.

