## Supplementary material for "Disruption of P2Y2 signaling promotes breast tumor cell dissemination by reducing ATP-dependent calcium elevation and actin localization to cell junctions": Combined Supplemental Figures and Legends

**SUPPLEMENTAL INFORMATION:**

**Supplemental Video 1. Representative full-length videos of breast epithelial cell response to ATP stimulation. A)** MCF10A cells loaded with Fluo-4 AM to label calcium and 10µM ATP or control was added after a 30 sec baseline read. Separate wells of cells treated with 10µM P2Y2i for 10 mins before imaging. Time stamp in bottom right corner in seconds with a frame rate of 20 fps.

**Supplemental Video 2. Representative full-length videos of mutant breast epithelial cells response to ATP stimulation. A)** 10A– PTEN^-/-^KRas cells loaded with Fluo-4 AM to label calcium and 10µM ATP or control was added after a 30 sec baseline read. Separate wells of cells treated with 10µM P2Y2i for 10 mins before imaging. Time stamp in bottom right corner in seconds with a frame rate of 20 fps.

**Supplemental Video 3. Representative full-length videos of metastatic breast cancer cells response to ATP stimulation. A)** MDA-MB-231 cells loaded with Fluo-4 AM to label calcium and 10µM ATP or control was added after a 30 sec baseline read. Separate wells of cells treated with 10µM P2Y2i for 10 mins before imaging. Time stamp in bottom right corner in seconds with a frame rate of 20 fps.

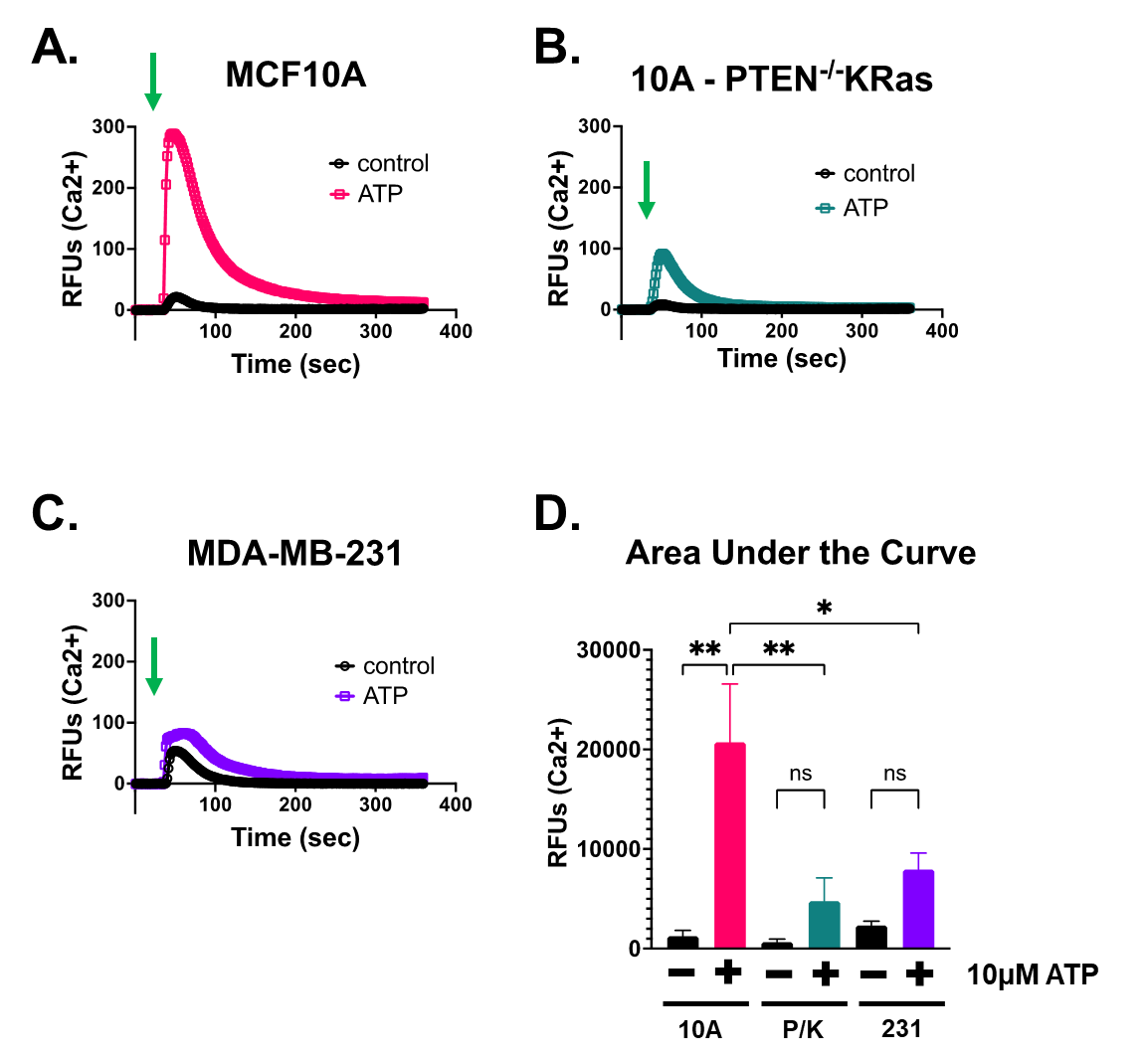

**Supplemental Figure 1. Quantitative measurements of intracellular calcium flux after ATP stimulation in non-malignant and metastatic breast epithelial cells.** Green arrows indicate when 10µM ATP was added. **A)** Quantification of calcium RFUs over time in MCF10A cells loaded with Fluo-4 AM. After a 30 second baseline read, 10µM ATP addition caused a rapid significant increase in Ca^2+^. **B)** The 10A-PTEN-/-KRas cells show a suppressed Ca^2+^ response. **C)** An inhibited, non-significant response is also seen in MDA-MB-231 cells after ATP stimulation. **D)** Area under the curve with one-way ANOVA and multiple comparisons was performed and there was a significant difference between control and ATP addition with SD (**, P<0.01, *, P<0.05) (n=3).

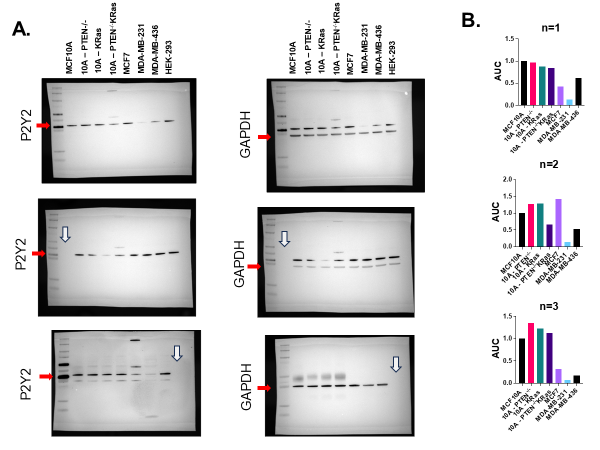

**Supplemental Figure 2. Protein expression of P2Y2 receptor across multiple breast epithelial cell lines. A)** Representative western blots from Figure 3. Western blots showed a decrease in P2Y2 expression in metastatic cell lines compared to MCF10A. GAPDH was used as a loading control and densitometry was performed. Red arrows indicate antibody. White arrows indicate an empty lane. **B)** Densitometry normalized to GAPDH and compared to MCF10A from all individual experiments.

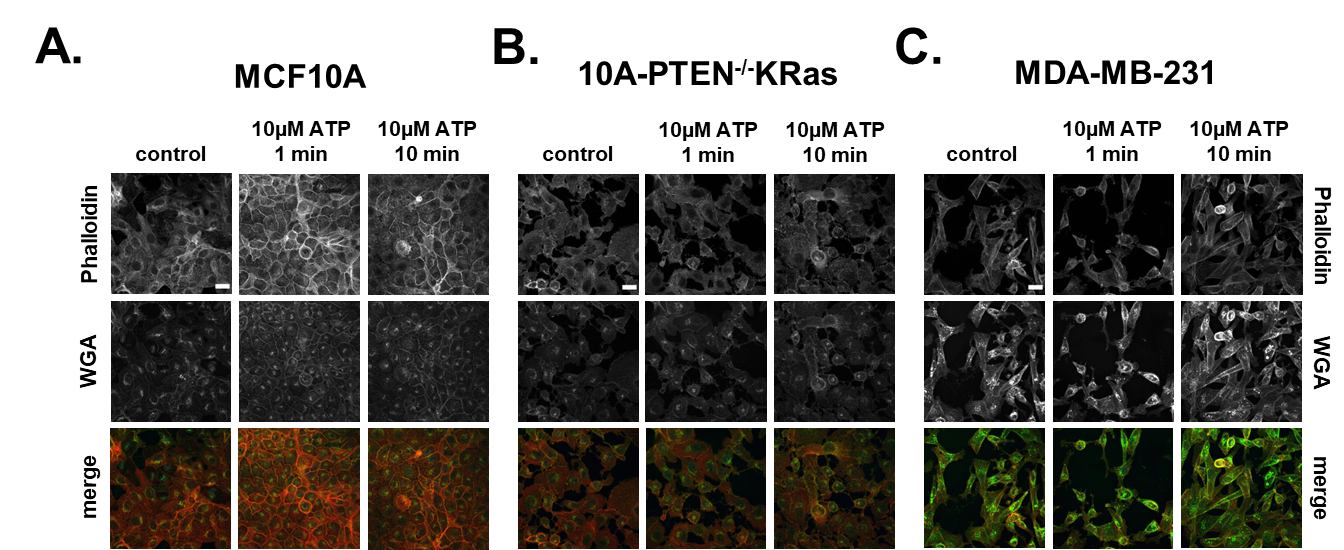

**Supplemental Figure 3. Immunofluorescence shows changes in actin in MCF10A cells while metastatic breast cancer cells have no changes.** All cells were treated with 10uM ATP for 1 min, 10 mins, or control. Representative images from Figure 5 stained with phalloidin, WGA, and DAPI. **A)** Confocal images of MCF10A cells showed actin polymerization and localization to cell edges and junctions. **B)** Confocal images of 10A-PTEN-/-KRas cells do not show major changes in phalloidin staining. **C)** Confocal images of MDA-MB-231 cells also show no changes in actin after ATP stimulation. All cells stained with Alexa conjugated Phalloidin (nm=647) and WGA (nm=488), while DAPI can be seen in the merged images stained with Hoechst (nm=461). Confocal NA: 60x/1.42 oil. Scale bar = 25µm.

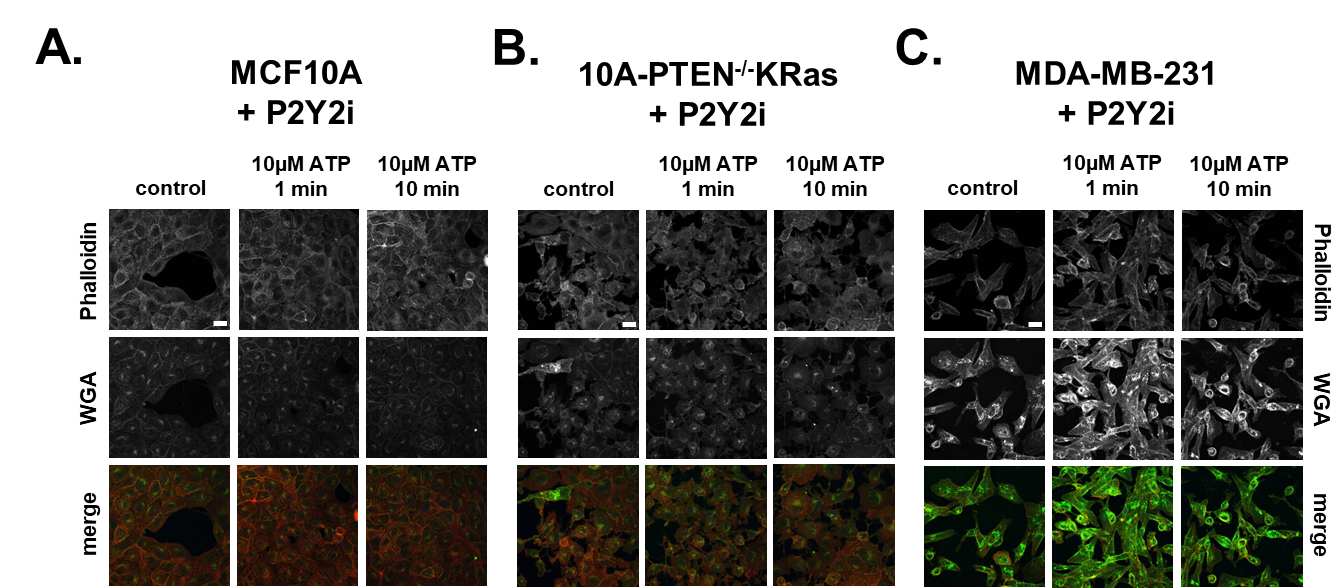

**Supplemental Figure 4. P2Y2i inhibits actin localization changes in non-tumorigenic breast epithelial cells.** All cells were pre-treated with 10uM P2Y2i for 10 mins, before ATP stimulation for 1 min, 10 mins, or control. Representative images from Figure 5 stained with phalloidin, WGA, and DAPI. **A.)** Confocal images of MCF10A cells treated with P2Y2i showed a lack of actin re-localization. **B.)** Confocal images of 10A-PTEN-/-KRas cells do not show major changes in phalloidin staining. **C.)** Confocal images of MDA-MB-231 cells also show no changes in actin after ATP stimulation. All cells stained with Alexa conjugated Phalloidin (nm=647) and WGA (nm=488), while DAPI can be seen in the merged images stained with Hoechst (nm=461). Confocal NA: 60x/1.42 oil. Scale bar = 25µm.

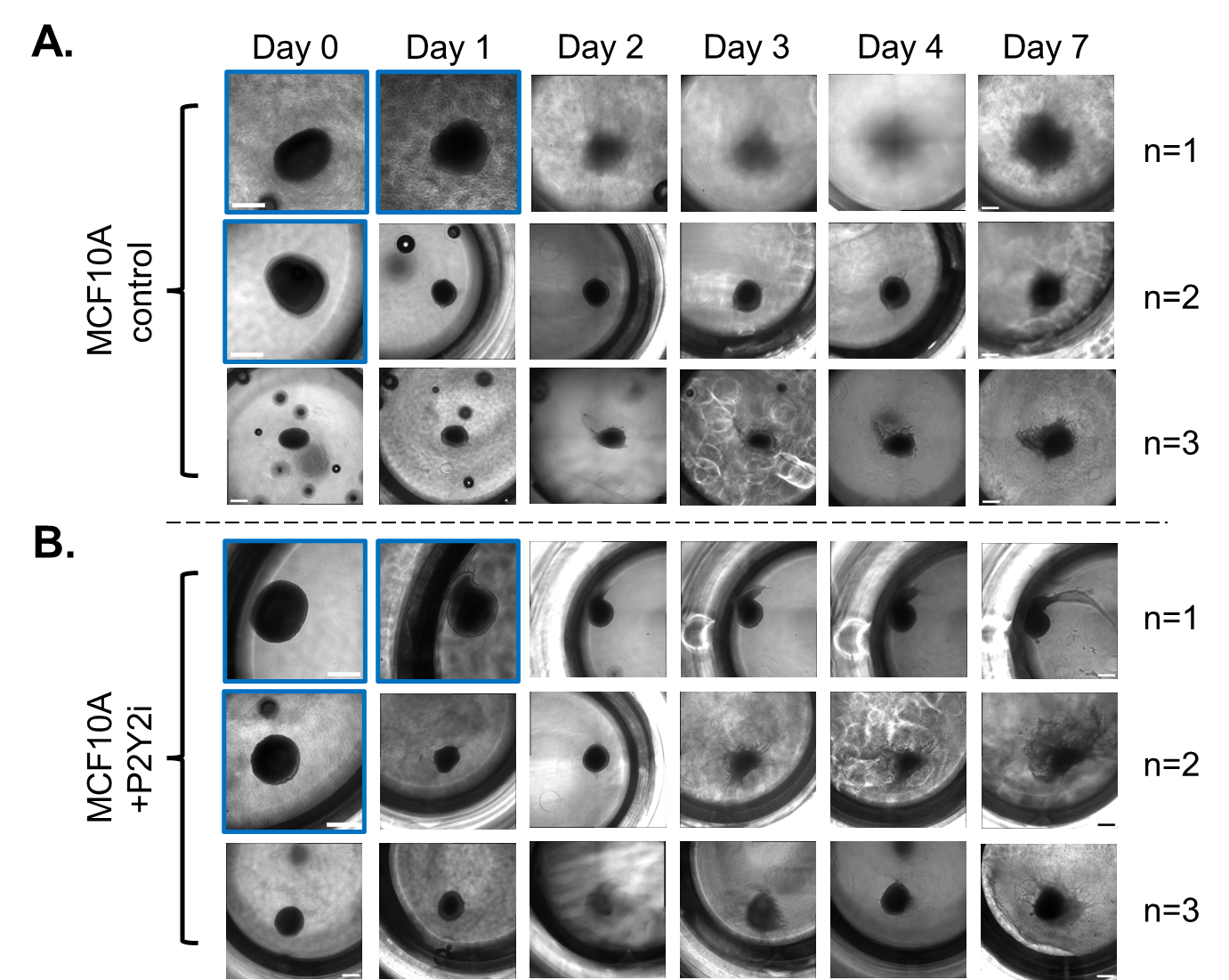

**Supplemental Figure 5. 3-D Cell Dissemination of non-tumorigenic MCF10A cells treated with P2Y2i.** Spheroids formed for 72hrs were embedded into collagen solution and imaged every day for 7 days. **A)** Representative phase contrast images taken at 10x on Day 0 (blue outline) or 10x stitched (2x2) of spheroids treated with control on Days 1, 4, and 7. **B)** Representative phase contrast images taken at 10x on Day 0 (blue outline) or 10x stitched (2x2) of spheroids treated with P2Y2i on Days 1, 4, and 7. Blue outlines represent 10x images unstitched (white or black scale bar=500µm).

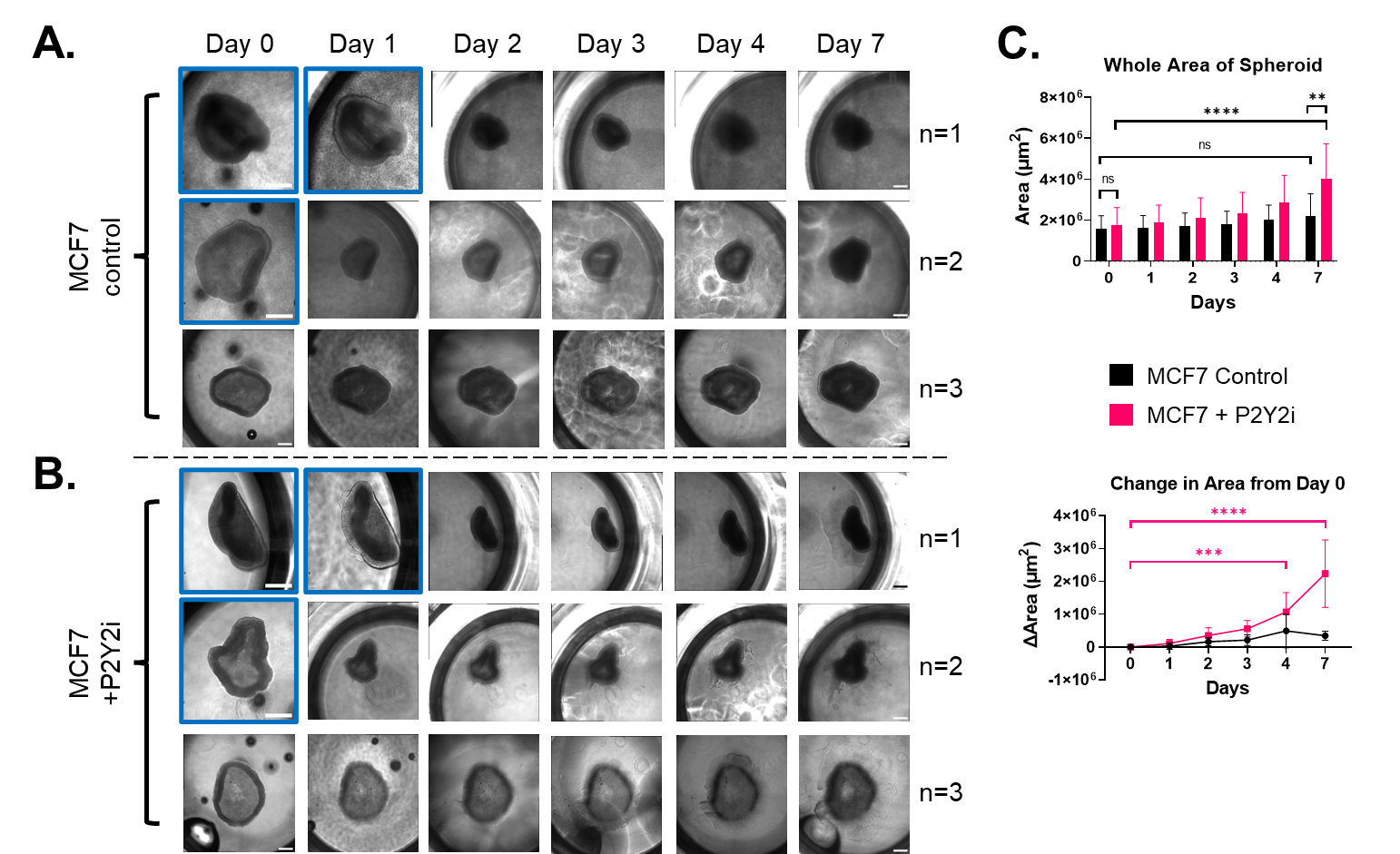

**Supplemental Figure 6. 3-D Cell Dissemination in tumorigenic, non-metastatic MCF7 cells treated with P2Y2i.** Spheroids formed for 72hrs were embedded into collagen solution and imaged every day for 7 days. **A)** Representative phase contrast images taken at 10x on Day 0 (blue outline) or 10x stitched (2x2) of spheroids treated with control on Days 1, 4, and 7. **B)** Representative phase contrast images taken at 10x on Day 0 (blue outline) or 10x stitched (2x2) of spheroids treated with P2Y2i on Days 1, 4, and 7. Blue outlines represent 10x images unstitched (white or black scale bar=500µm). **C)** Graphs showing whole area of spheroid (left) and change in area (µm^2^) from day 0 through day 7 (right) with SD, and two-way ANOVA performed to calculate significance (****, P<0.0001, ***, P<0.0005, **, P<0.005).
